# B-cell receptor reconstruction from single-cell RNA-seq with VDJPuzzle

**DOI:** 10.1101/181156

**Authors:** Simone Rizzetto, David NP Koppstein, Jerome Samir, Mandeep Singh, Joanne H. Reed, Curtis H. Cai, Andrew R. Lloyd, Auda A. Eltahla, Christopher C. Goodnow, Fabio Luciani

## Abstract

The B-cell receptor (BCR) performs essential functions for the adaptive immune system including recognition of pathogen-derived antigens. Cell-to-cell variability of BCR sequences due to V(D)J recombination and somatic hypermutation (SHM) necessitates single-cell characterization of BCR sequences. Single-cell RNA sequencing (scRNA-seq) presents the opportunity for simultaneous capture of the BCR sequence and transcriptomic signature for a detailed understanding of the dynamics of an immune response.

We developed VDJPuzzle 2.0, a bioinformatic tool that reconstructs productive, full-length B-cell receptor sequences of both heavy and light chains. VDJPuzzle successfully reconstructs BCRs from 98.3% (n=117) of human and 96.5% (n=200) from murine B cells. 92.0% of clonotypes and 90.3% of mutations were concordant with single-cell Sanger sequencing of the immunoglobulin chains. VDJPuzzle is available at https://bitbucket.org/kirbyvisp/vdjpuzzle2

## Introduction

B cells are an essential component of the adaptive immune system. Specificity of B cells for their antigen is imparted by the BCR, which comprises a paired heavy (IGH) and light (IGK or IGL) immunoglobulin chain that differs from cell to cell. Each of these chains contains one of many possible V, J, and constant segments encoded by clusters of genomically-adjacent genes. Additionally, IGH includes a D segment between the V and J segments. The specific combination of genes is selected through genomic rearrangement in a process called V(D)J recombination and occurs during B-cell development [1].

After V(D)J recombination, naïve B cells are activated by binding an antigen through the BCR. Additional BCR diversity and specificity for antigen is generated through a process called affinity maturation during which SHMs accumulate in the germline sequence of the V(D)J genes. Preferential mutation occurs in the complementary-determining regions (CDRs) which directly bind to antigen; the most variable of these is the complementarity determining region 3 (CDR3) which adjoins the V(D)J junction. This region determines the affinity of the BCR for the cognate antigen.

Obtaining high-quality BCR sequences along with single-cell transcriptomics is crucial for characterizing the dynamics of B-cell evolution and differentiation during an immune response. scRNA-seq enables the simultaneous measurement of these important parameters [2, 3]. Similar approaches for reconstruction of full-length T-cell receptors (TCRs) have yielded insights into T cell biology [4-8], including determination of T-cell development trajectories in the tumor microenvironment [9].

Here, we present VDJPuzzle 2.0, a computational method to reconstruct the full length BCRs from scRNA-seq and at the same time also provide gene expression data. This tool extends a software package previously developed to reconstruct T-cell receptors [6]. The new VDJPuzzle version has been adapted for BCRs and includes additional features to reliably characterise SHMs.

## Results

### Algorithm

For each cell, trimmed paired-end reads are aligned to the reference genome. Pairs with at least one read aligning to any of the V(D)J or constant-region genes are extracted and used to assemble contigs using the *de novo* assembler Trinity [10]. Assembled contigs are matched to the IMGT database using MiGMAP (https://github.com/mikessh/migmap) to find complete, productive, and in-frame BCRs [11]. VDJPuzzle workflow is represented in Fig. 1.

**Fig. 1.**
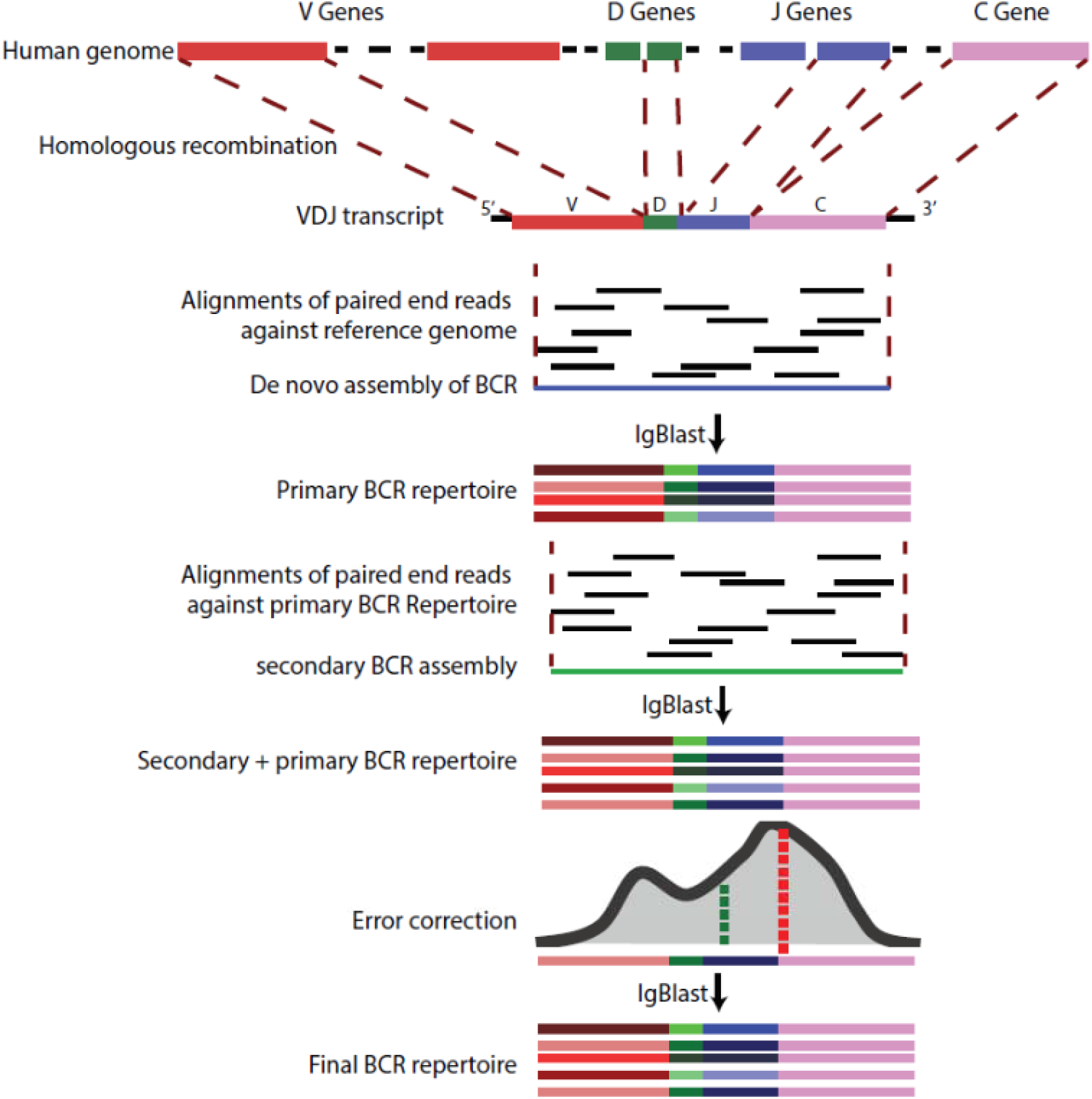
VDJPuzzle workflow.

### Success rate and quality of reconstructed BCR sequences

VDJPuzzle was applied to scRNA-seq data from 117 B cells isolated from human peripheral blood mononuclear cells (PBMCs) derived from a healthy donor. These B cells were additionally sorted by surface phenotype: 27 transitional, 30 naïve, 30 memory, and 30 plasmablast B cells. We define a successful BCR reconstruction as a chain with a complete, productive, and in-frame BCR sequence identified using IgBLAST run via MiGMAP. VDJPuzzle successfully reconstructed at least one V(D)J sequence of a heavy chain (IGH) and at least one light chain (IGL or IGK) in 98.3% (n=115). Of the 115 B cells, 72 (62.6%) contained at least one IGK, while 54 (47.0%) had at least one IGL (Table 1). In 7.0% (n=8) cells, VDJPuzzle reconstructed both IGK and IGL, consistent with previous findings of isotypic inclusion within the normal B cell repertoire of hu-mans and mice [12].

**Table 1.** Benchmark results of the VDJPuzzle, BASIC, and Sanger sequencing.

| Method | IGH | IGK | IGL |
| --- | --- | --- | --- |
| VDJPuzzle | 115 (98%) | 72 (61.5%) | 54 (45.3%) |
| BASIC | 7 (6.0%) | 19 (16.3%) | 6 (5.1%) |
| Sanger | 95 (81.2%) | 55 (47%) | 25 (21.4%) |

Sanger sequencing from the single-cell cDNA libraries for the 117 B cells was performed as validation of the VDJPuzzle output. After filtering low-quality traces, Sanger sequencing reconstructed BCR transcripts in 95 and 80 cells, respectively. Out of the 95 IGH, 89 had concordant clonotypes (i.e. identical CDR3 and V-J genes) compared to the corresponding VDJPuzzle-reconstructed BCR. Two cells had the same CDR3 sequence, but discordant V and J genes. Two other cells had both different V-J genes and CDR3 sequences. VDJPuzzle failed to reconstruct the receptor in two other cells. A total of 78 light chain (24 IGL and 52 IGK) sequences from the 80 reconstructed with Sanger matched the VDJPuzzle result in both CDR3 and V-J genes. The two discordant chains contained either a single amino acid difference in the CDR3 sequence, or had the same CDR3 but different V-J genes.

Analysis of mutations with respect to the germline from the 89 heavy chains and 78 light chains with concordant clonotypes between Sanger and VDJPuzzle revealed that 2376 (97.7%) mutations in the heavy chain and 1666 (94.9%) in the light chain were shared with Sanger. In contrast, 57 and 89 were identified only by VDJpuzzle, while 141 and 226 mutations were identified by Sanger alone.

As further comparison, a recently-published BCR reconstruction method called BASIC was tested on the same dataset [13]. Only 6.0% (n=7) of heavy chains and 21.4% (n=25) of light chains were complete, productive, and in-frame as determined by IgBLAST [14]. However, BASIC success rate may improve with a different dataset or when a different definition of successful reconstructed BCR is used. Indeed, Canzar et al. reported a success rate of 97% in identifying the correct V(D)J gene usage.

As alternative comparison, a less stringent success rate has been utilized to benchmark VDJPuzzle, BASIC [13], and Sanger sequencing. In this analysis, we allowed MiGMAP to report BCR with incomplete, not in-frame, non-coding, and not canonical BCRs. Results are showed in Table 2. Both VDJPuzzle and BASIC improved, however this also increased the number of cells with multiple heavy and light chains due to the inclusion of non-productive BCR.

**Table 2.** Alternative benchmarking. Numbers and percentage of reconstructed chains considering only BCRs with at least one V and J (no D and CDR3 required).

| Method | IGH | IGK | IGL |
| --- | --- | --- | --- |
| VDJPuzzle | 117 (100%) | 104 (88.9%) | 103 (88.0%) |
| BASIC | 59 (50.4%) | 59 (50.4%) | 55 (47.0%) |
| Sanger | 96 (82.1%) | 55 (47%) | 26 (22.2%) |

Next, we examined germline mutations as a function of IMGT-annotated framework (FR) and CDR regions [11]. As expected, more SHMs occurred in the CDR regions compared to the FR regions. Further, plasmablasts and memory cells carried more mutations, while naïve and transitional B cells had a lower number of SHMs (Fig. 2). To demonstrate the utility of the VDJPuzzle output, we linked the aggregate mutation rate across the entire V(D)J region with the gene-expression data from scRNA-seq (Fig. 3). Unsupervised clustering of the transcriptomic data identified markers of naïve/transitional, plasmablast, and memory populations. Known terminal differentiation markers such as XBP1 were highly expressed in the plasmablast cluster [15], whereas expression of known early differentiation markers including TCL1A was observed in naïve/transitional cells [16]. Furthermore, a cluster of co-expressed genes was negatively correlated with mutation frequencies (R=-0.79, p-value<0.001).

**Fig. 2.**
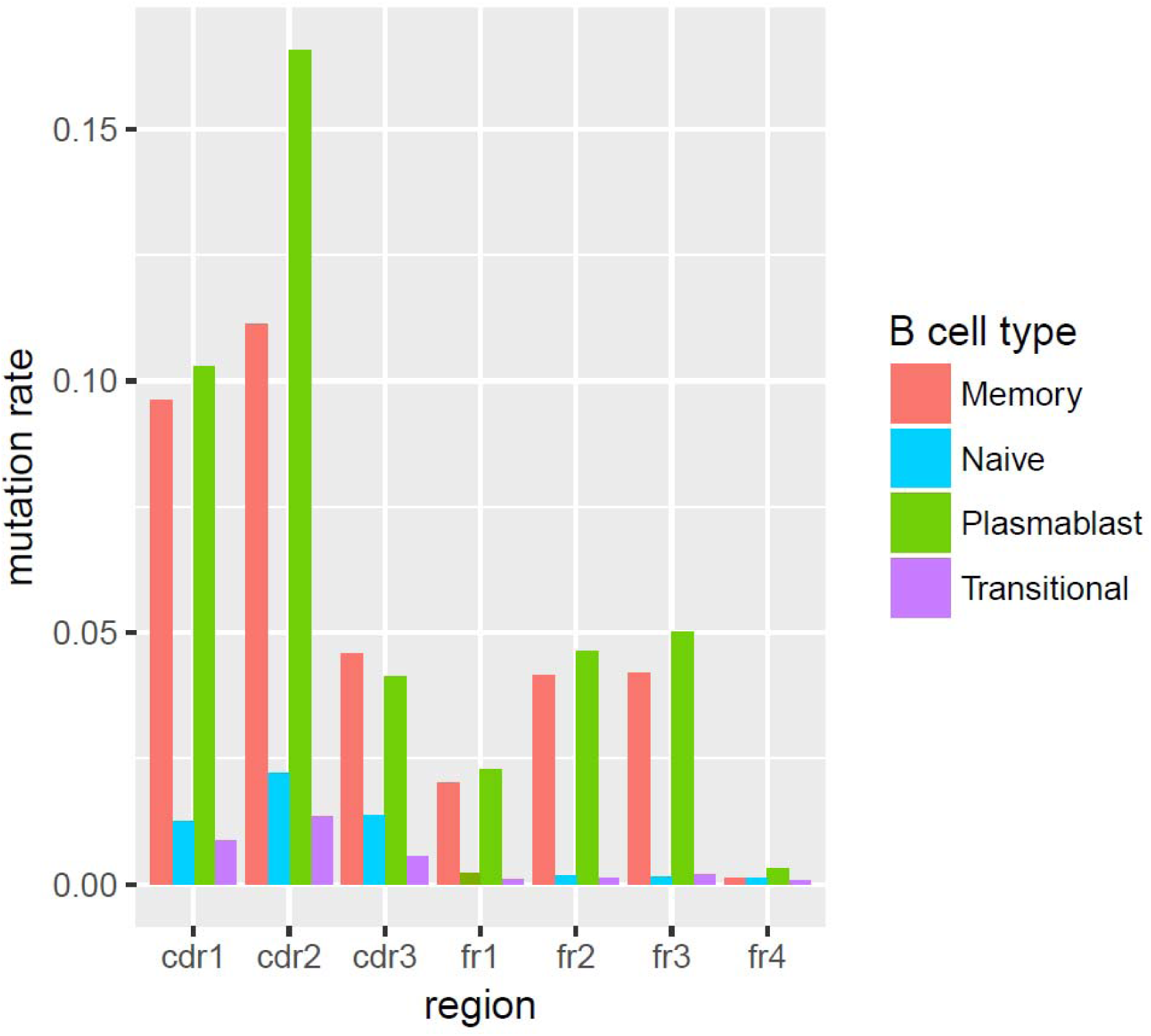
Mutation rate across FR and CDR regions. Mutations are called comparing the sequence against the BCR germline sequence. The number of mutations is normalized by the length of the FR/CDR regions.

**Fig. 3.**
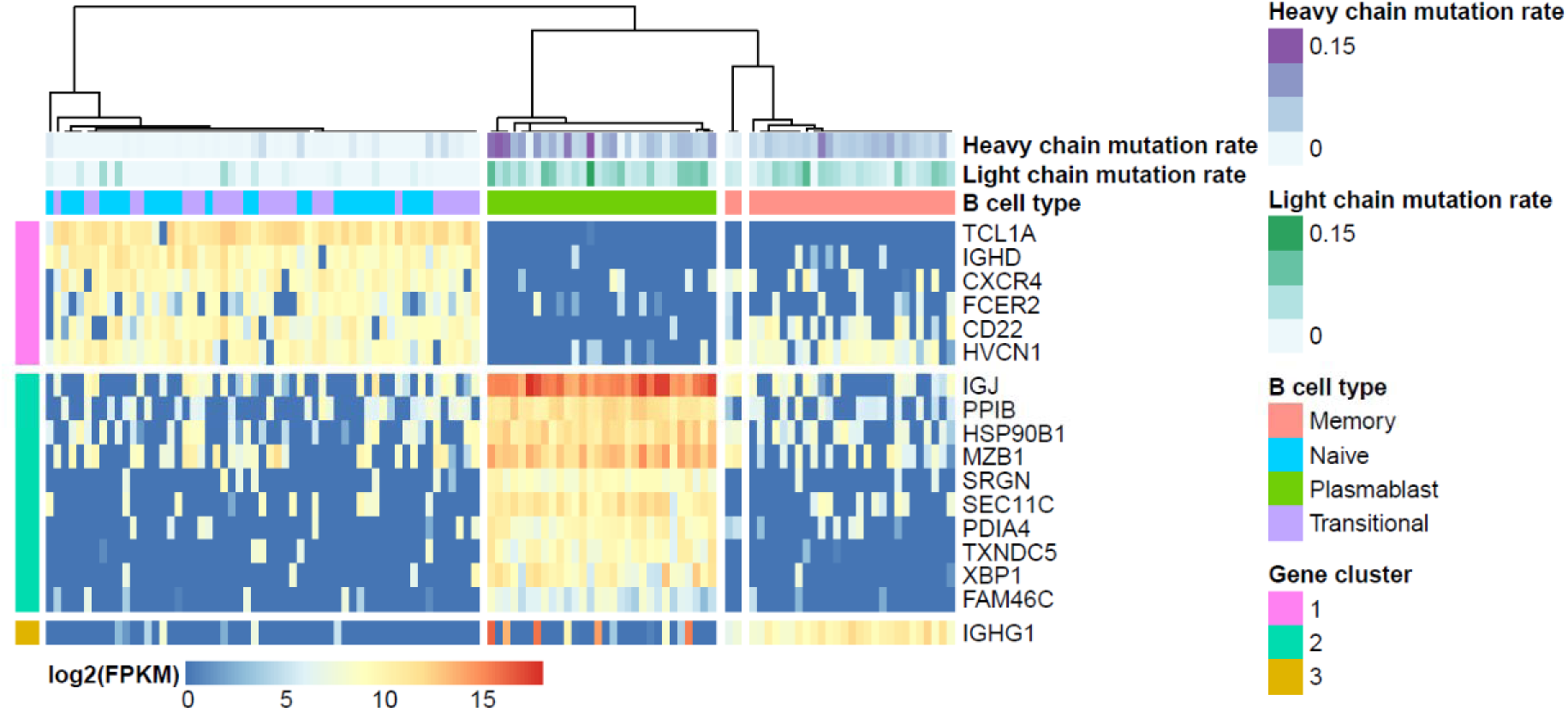
Clustering analysis of scRNAseq and relationship with SHMs. Unsupervised clustering was performed using SC3. Naive and transient B cells are clustered together and associated to a low mutation rate. In contrast, Memory B cells and Plasmablast form two clusters which are associated to high mutation rate. Notably, the gene signature (gene cluster 1) associated to naïve and transient B cells was negatively correlated to the average mutation rate identified in each cell.

VDJPuzzle was also tested on mouse B cells. Out of 213 cells in ArrayExpress (E-MTAB-4825), 13 were not available for download on the FTP server. From the remaining 200 cells, VDJPuzzle reconstructed 193 heavy chains (96.5%) and 200 light chains (100%). 195 cells (97.5%) had at least one IGK, while 8 (4%) had at least one IGL. 3 cells (1.5%) had both IGK and IGL.

## Discussion

VDJPuzzle achieves a high success rate in reconstructing BCR sequence from scRNA-seq compared to existing methods. Other software can achieve high detection of V(D)J usage and the CDR3 from RNA-seq data, such as MiXCR [17]; however, these programs cannot reconstruct the full-length BCR sequence and do not integrate these data with gene expression. We anticipate that simultaneous determination of BCR sequences with transcriptomic signatures from single cells will be a promising approach for studying the dynamics of an immune response in heterogeneous populations of B cells.

## Methods

### Single cell sorting

PBMCs were separated from freshly collected whole blood by density gradient centrifugation and stained with the following monoclonal antibodies: CD19-BV421 (Biolegend), IgD-PerCP-Cy5.5, CD27-PE-Cy7, CD38-APC, CD10-PECF594, CD3-APC-Cy7, IgG-BV605 (BD Biosciences) and viability dye eFluor780 (eBioscience) for 30 min on ice and, following two washes, cells were resuspended in PBS with 1% HI bovine serum for single cell sorting. Following exclusion of non-viable/CD3+ cells, individual transitional (CD19+, IgD+, CD27-, CD10+), mature naïve (CD19+, IgD+, CD27-, CD10-), switched memory (CD19+, IgD-, CD27+, IgG+) and plasmablast (CD19+, IgD-, CD27++, CD38++) were sorted directly into 96-well PCR plates (Eppendorf) using a FACS ARIA III (BD Biosciences). PCR plates were stored at –80 C for subsequent analysis.

### Single cell RNA-sequencing protocol

Single cell RNA sequencing was performed as described by Picelli et al. [18, 19] with the following modifications. Cell lysis buffer was prepared by adding 1μL Rnase inhibitor (Clontech, Mountain View, CA) to 19μl Triton X-100 solution (0.2% v/v). Cells were sorted directly into 96-well plates containing 1μL lysis buffer, 0.5μL dNTP (10mM) and 0.5μL oligo-dT primer at 5uM. Reverse-transcription and PCR amplification were performed as described with the following exceptions: reactions were performed at half volumes, the IS PCR primer was reduced to a 50nM final concentration and the number of PCR cycles increased to 28. Sequencing libraries were prepared using the Nextera XT Library Preparation Kit (Illumina; San Diego, CA, USA) at one quarter of the recommend total volume and sequencing performed on the Illumina NextSeq sequencing platform.

### Single cell RNA-sequencing data analysis

Reads obtained from scRNA-seq experiments were used as input of VDJPuzzle which perform the BCR assembly (method section of the main text), it aligns the reads to the ensembl reference genome (GRCh37), and it quantifies the gene expression of each cell. VDJPuzzle alignment step utilized TopHat2 [20], while gene expression is calculated with CuffLinks 2.2.1 [21]. Downstream analysis of gene expression has been performed in R. Clustering analysis have been performed with SC3 R package [22].

### Sanger sequencing protocol

Sanger sequencing was performed fromorm the same cDNA libraries utilized for scRNAseq for the 117 B cells and for both heavy and light chains. Two ng of amplified SmartSeq2 cDNA was subjected to 35 cycles of PCR using forward primers binding leader peptide sequences of immunoglobulin heavy and light chain V segments and reverse primers binding μ, γ, κ, and λ constant regions [23]. PCR products were Sanger sequenced with the same forward and reverse primers by Genewiz (Boston, USA).

### Sanger sequences data analysis

The 468 tracing files (forward/reverse for each chain and each cell) were loaded on Geneious [24] and manually trimmed based on the chromatogram quality. Forward and reverse sequences were merged together to create a consensus sequence for each chain and each cell. The sequences were loaded into MiGMAP to identify the V(D)J usages, FRs and CDRs regions, along with the mutations compared to the germline sequence of the matched BCR in IMGT.

### Single cell RNA-sequencing of mouse B cells

Mouse scRNA-seq were downloaded from ArrayExpress (E-MTAB-4825) [3]. Data analysis was performed as for human PBMC using GRCm38 reference genome on the 200 B cells retrieved from the ArrayExpress ftp server.

## References

1. Tonegawa S. Somatic generation of antibody diversity. Nature. 1983;302(5909):575–81. doi: 10.1038/302575a0.

2. Lizotte PH, Ivanova EV, Awad MM, Jones RE, Keogh L, Liu H, et al. Multiparametric profiling of non-small-cell lung cancers reveals distinct immunophenotypes. JCI Insight. 2016;1(14):e89014. doi: 10.1172/jci.insight.89014. PubMed PMID: 27699239; PubMed Central PMCID: PMCPMC5033841.

3. Wu YL, Stubbington MJ, Daly M, Teichmann SA, Rada C. Intrinsic transcriptional heterogeneity in B cells controls early class switching to IgE. J Exp Med. 2017;214(1):183–96. doi: 10.1084/jem.20161056. PubMed PMID: 27994069; PubMed Central PMCID: PMCPMC5206502.

4. Rizzetto S, Eltahla AA, Lin P, Bull R, Lloyd A, Ho JWK, et al. Impact Of Sequencing Depth And Read Length On Single Cell RNA Sequencing Data: Lessons From T Cells. bioRxiv. 2017. doi: 10.1101/134130.

5. Redmond D, Poran A, Elemento O. Single-cell TCRseq: paired recovery of entire T-cell alpha and beta chain transcripts in T-cell receptors from single-cell RNAseq. Genome Med. 2016;8(1):80. doi: 10.1186/s13073-016-0335-7. PubMed PMID: 27460926; PubMed Central PMCID: PMCPMC4962388.

6. Eltahla AA, Rizzetto S, Rasoli M, Betz-Stablein BD, Venturi V, Kedzierska K, et al. Linking the T cell receptor to the single cell transcriptome in antigen-specific human T cells. Immunology and cell biology. 2016.

7. Stubbington MJT, Lonnberg T, Proserpio V, Clare S, Speak AO, Dougan G, et al. T cell fate and clonality inference from single-cell transcriptomes. Nat Methods. 2016;13(4):329–32. doi: 10.1038/nmeth.3800. PubMed PMID: 26950746; PubMed Central PMCID: PMCPMC4835021.

8. Afik S, Yates KB, Bi K, Darko S, Godec J, Gerdemann U, et al. Targeted reconstruction of T cell receptor sequence from single cell RNA-seq links CDR3 length to T cell differentiation state. Nucleic acids research. 2017. doi: 10.1093/nar/gkx615.

9. Zheng C, Zheng L, Yoo JK, Guo H, Zhang Y, Guo X, et al. Landscape of Infiltrating T Cells in Liver Cancer Revealed by Single-Cell Sequencing. Cell. 2017;169(7):1342–56 e16. doi: 10.1016/j.cell.2017.05.035. PubMed PMID: 28622514.

10. Grabherr MG, Haas BJ, Yassour M, Levin JZ, Thompson DA, Amit I, et al. Full-length transcriptome assembly from RNA-Seq data without a reference genome. Nature biotechnology. 2011;29(7):644–52. doi: 10.1038/nbt.1883. PubMed PMID: 21572440; PubMed Central PMCID: PMCPMC3571712.

11. Lefranc MP, Giudicelli V, Duroux P, Jabado-Michaloud J, Folch G, Aouinti S, et al. IMGT(R), the international ImMunoGeneTics information system(R) 25 years on. Nucleic acids research. 2015;43(Database issue):D413–22. doi: 10.1093/nar/gku1056. PubMed PMID: 25378316; PubMed Central PMCID: PMCPMC4383898.

12. Giachino C, Padovan E, Lanzavecchia A. kappa+lambda+ dual receptor B cells are present in the human peripheral repertoire. J Exp Med. 1995;181(3):1245–50. PubMed PMID: 7869042; PubMed Central PMCID: PMCPMC2191910.

13. Canzar S, Neu KE, Tang Q, Wilson PC, Khan AA. BASIC: BCR assembly from single cells. Bioinformatics. 2017;33(3):425–7. doi: 10.1093/bioinformatics/btw631. PubMed PMID: 28172415; PubMed Central PMCID: PMCPMC5408917.

14. Ye J, Ma N, Madden TL, Ostell JM. IgBLAST: an immunoglobulin variable domain sequence analysis tool. Nucleic acids research. 2013;41(Web Server issue):W34–40. doi: 10.1093/nar/gkt382. PubMed PMID: 23671333; PubMed Central PMCID: PMCPMC3692102.

15. Reimold AM, Iwakoshi NN, Manis J, Vallabhajosyula P, Szomolanyi-Tsuda E, Gravallese EM, et al. Plasma cell differentiation requires the transcription factor XBP-1. Nature. 2001;412(6844):300–7. doi: 10.1038/35085509. PubMed PMID: 11460154.

16. Teitell MA. The TCL1 family of oncoproteins: co-activators of transformation. Nat Rev Cancer. 2005;5(8):640–8. doi: 10.1038/nrc1672. PubMed PMID: 16056259.

17. Bolotin DA, Poslavsky S, Mitrophanov I, Shugay M, Mamedov IZ, Putintseva EV, et al. MiXCR: software for comprehensive adaptive immunity profiling. Nat Methods. 2015;12(5):380–1. doi: 10.1038/nmeth.3364. PubMed PMID: 25924071.

18. Picelli S, Faridani OR, Björklund AK, Winberg G, Sagasser S, Sandberg R. Full-length RNA-seq from single cells using Smart-seq2. Nature protocols. 2014;9(1):171–81. doi: 10.1038/nprot.2014.006.

19. Picelli S, Björklund ÅKK, Faridani OR, Sagasser S, Winberg G, Sandberg R. Smart-seq2 for sensitive full-length transcriptome profiling in single cells. Nature methods. 2013;10(11):1096–8. doi: 10.1038/nmeth.2639.

20. Kim D, Pertea G, Trapnell C, Pimentel H, Kelley R, Salzberg SL. TopHat2: accurate alignment of transcriptomes in the presence of insertions, deletions and gene fusions. Genome Biol. 2013;14(4):R36. doi: 10.1186/gb-2013-14-4-r36. PubMed PMID: 23618408; PubMed Central PMCID: PMCPMC4053844.

21. Trapnell C, Williams BA, Pertea G, Mortazavi A, Kwan G, van Baren MJ, et al. Transcript assembly and quantification by RNA-Seq reveals unannotated transcripts and isoform switching during cell differentiation. Nature biotechnology. 2010;28(5):511–5. doi: 10.1038/nbt.1621. PubMed PMID: 20436464; PubMed Central PMCID: PMCPMC3146043.

22. Kiselev VY, Kirschner K, Schaub MT, Andrews T, Yiu A, Chandra T, et al. SC3: consensus clustering of single-cell RNA-seq data. Nat Methods. 2017;14(5):483–6. doi: 10.1038/nmeth.4236. PubMed PMID: 28346451; PubMed Central PMCID: PMCPMC5410170.

23. Tiller T, Meffre E, Yurasov S, Tsuiji M, Nussenzweig MC, Wardemann H. Efficient generation of monoclonal antibodies from single human B cells by single cell RT-PCR and expression vector cloning. J Immunol Methods. 2008;329(1-2):112–24. doi: 10.1016/j.jim.2007.09.017. PubMed PMID: 17996249; PubMed Central PMCID: PMCPMC2243222.

24. Kearse M, Moir R, Wilson A, Stones-Havas S, Cheung M, Sturrock S, et al. Geneious Basic: an integrated and extendable desktop software platform for the organization and analysis of sequence data. Bioinformatics. 2012;28(12):1647–9. doi: 10.1093/bioinformatics/bts199. PubMed PMID: 22543367; PubMed Central PMCID: PMCPMC3371832.

